# Mindfulness-based Neurofeedback: A Systematic Review of EEG and fMRI studies

**DOI:** 10.1101/2024.09.12.612669

**Authors:** Isaac N. Treves, Keara D. Greene, Zia Bajwa, Emma Wool, Nayoung Kim, Clemens C.C. Bauer, Paul A. Bloom, David Pagliaccio, Jiahe Zhang, Susan Whitfield-Gabrieli, Randy P. Auerbach

## Abstract

Neurofeedback concurrent with mindfulness meditation may reveal meditation effects on the brain and facilitate improved mental health outcomes. Here, we systematically reviewed EEG and fMRI studies of mindfulness meditation with neurofeedback (mbNF) and followed PRISMA guidelines. We identified 10 fMRI reports, consisting of 177 unique participants, and 9 EEG reports, consisting of 242 participants. Studies of fMRI focused primarily on downregulating the default-mode network (DMN). Although studies found decreases in DMN activations during neurofeedback, there is a lack of evidence for transfer effects, and the majority of studies did not employ adequate controls, e.g. sham neurofeedback. Accordingly, DMN decreases may have been confounded by general task-related deactivation. EEG studies typically examined alpha, gamma, and theta frequency bands, with the most robust evidence supporting the modulation of theta band activity. Both EEG and fMRI mbNF have been implemented with high fidelity in clinical populations. However, the mental health benefits of mbNF have not been established. In general, mbNF studies would benefit from sham-controlled RCTs, as well as clear reporting (e.g. CRED-NF).

## 1 Introduction

Mindfulness meditation involves cultivating an accepting, open-minded attention to the present moment (Creswell, 2017). The word mindfulness originated from Eastern contemplative traditions, specifically, as a translation of the term *sati* from Pali or *smrti* from Sanskrit, which mean remembering or being aware. Mindfulness was largely introduced to Western medicine with the advent of mindfulness-based stress reduction (MBSR) in the 1980s (Kabat-Zinn, 1982, 2003). MBSR and its adaptations have been used to treat chronic pain (Goyal et al., 2014; Kabat-Zinn, 1982), anxiety (Goldin et al., 2013; Hoge et al., 2022; Hölzel et al., 2013), addiction (Black, 2014; Garland et al., 2015; Vallejo & Amaro, 2009), and depression (Kuyken et al., 2016). Indeed, mindfulness has been documented to be equally effective as pharmacological treatment for anxiety disorders (Hoge et al., 2022) and potentially more effective than cognitive behavioral therapy (CBT) for treatment of mild-to-moderate depression (Strauss et al., 2023). Mindfulness is now a central component of leading psychotherapeutic approaches like dialectical behavioral therapy (DBT) (Linehan, 1993; McCauley et al., 2018) and acceptance and commitment therapy (ACT) (Hayes et al., 1999).

Mindfulness meditations include practices like *breath awareness*, which involves orienting attention to one’s breath and practicing returning to the breath every time one’s attention wanders away, and *body scans*, involving moving the spotlight of attention from body part to body part with a curious and non-judgmental attitude towards the sensations one encounters. Another practice is *open monitoring*, where one notices transient thoughts and sensations in an open state without attaching to them. Breath awareness and body scans are often called *focused attention* (FA) practices, aiming to cultivate a stable and precise attention, which contrasts with *open monitoring* (OM) practices, cultivating receptivity to experience (Lutz et al., 2008).

There are several theories regarding the neurobiological mechanisms behind mindfulness meditation. One influential account suggests that large-scale brain networks are involved (Hasenkamp et al., 2012; Mooneyham et al., 2016). Specifically, this account implicates the default-mode network (DMN), with core regions of the posterior cingulate cortex (PCC) and medial prefrontal cortex (mPFC) as well as the central executive network (CEN), with core regions of the dorsolateral prefrontal cortex (DLPFC) and parietal cortex, and the salience network (SN), with core regions of the anterior cingulate cortex (ACC) and insula **(Figure 1**). In this account, the DMN is involved in mind-wandering away from the object of meditation, the CEN is involved in goal-directed maintenance of the object, and the SN is involved in switching between the two (Hasenkamp et al., 2012). This is based largely on functional magnetic resonance imaging (fMRI) during focused attention meditation (Fox et al., 2016; Ganesan et al., 2022), changes observed in networks after mindfulness training (Rahrig et al., 2022; Sezer et al., 2022), in addition to a robust cognitive neuroscience literature on these networks (Menon, 2011). In tandem, researchers have examined changes in brain rhythms or oscillations during meditation using electroencephalography (EEG) (**Figure 2**). Brain oscillations represent information processing across wide-ranging brain regions, and change with attention (Herrmann et al., 2016). There is evidence of power increases in alpha, theta and gamma waves during meditation (Chiesa & Serretti, 2010; Lee et al., 2018; Lomas et al., 2015; Stapleton et al., 2020). Alpha and theta power may correspond to inwardly focused attention (Lomas et al., 2015), whereas gamma power may reflect broad awareness (Lomas et al., 2015; Lutz et al., 2004; Stapleton et al., 2020). Despite this meaningful work, the field still lacks a complete mechanistic account of mindfulness meditation. Take, for example, the assertion that mindfulness decreases DMN activation (Brewer et al., 2011; Ganesan et al., 2022). This particular assertion is largely founded on comparisons of meditation to control conditions, which do not directly imply mechanistic involvement. For example, neural changes associated with mindfulness may be caused by decreases in stress accompanying meditation, rather than the voluntary and directed actions of meditation. In addition, the choice of control condition can lead to differing results. For example, Ganesan and colleagues (2022), found that the DMN was less activated during meditation than control conditions in only 60% of the studies reviewed, with the controls including rest, intentional instructions to mind wander, and other functional tasks. A final concern is that reverse inferences from brain areas to psychological processes may be implausible (Poldrack, 2006). To test theories about mindfulness meditation and uncover a more complete mechanistic account, researchers need to manipulate brain function, and neurofeedback affords one opportunity to manipulate brain functions directly implicated in mindfulness meditation.

**Figure 1:**
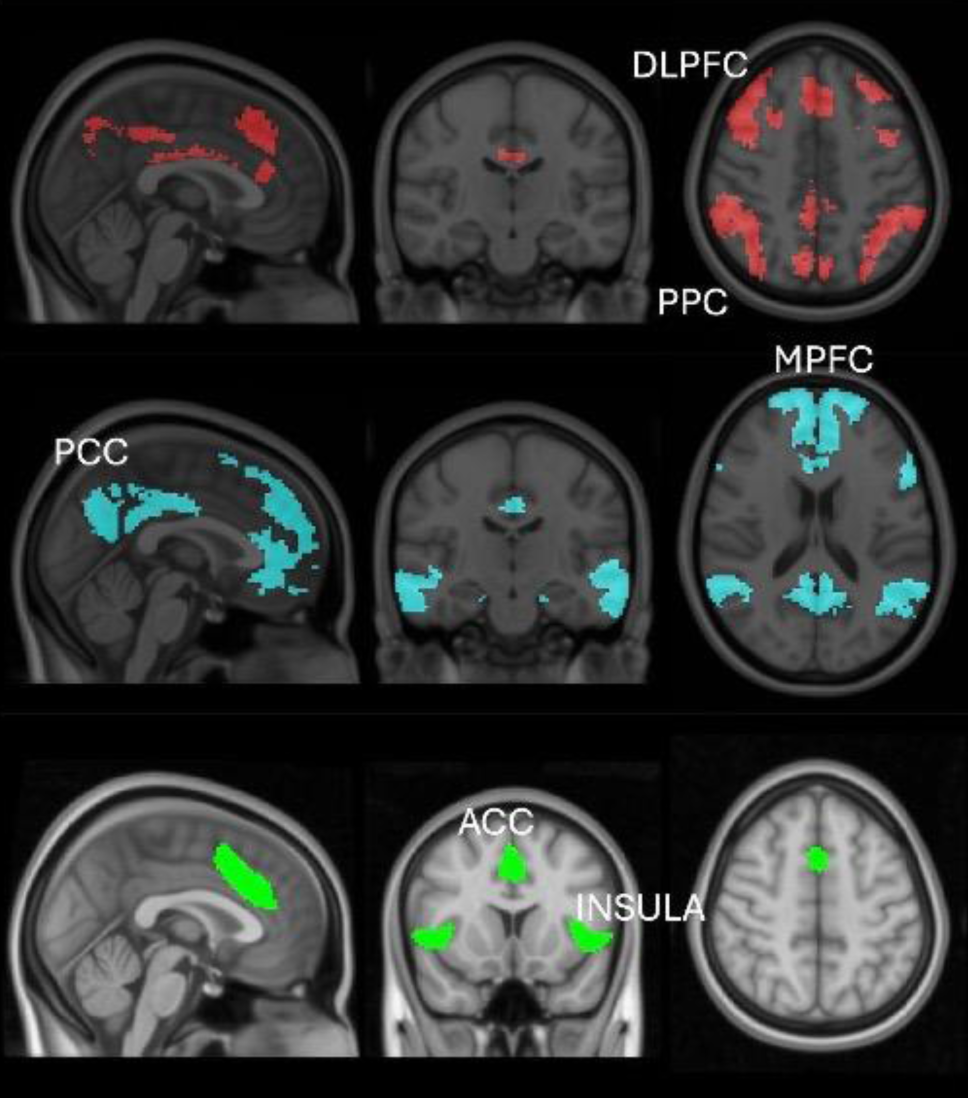
Brain networks involved in mindfulness meditation. Central executive network, in red; Default-mode network, in blue; Salience network, in green. DLPFC: dorsolateral prefrontal cortex; PPC: posterior parietal cortex; PCC: posterior cingulate cortex; MPFC: medial prefrontal cortex; ACC: anterior cingulate cortex; Insula: insular cortex. Adapted with permission from Treves et al., in press.

**Figure 2:**
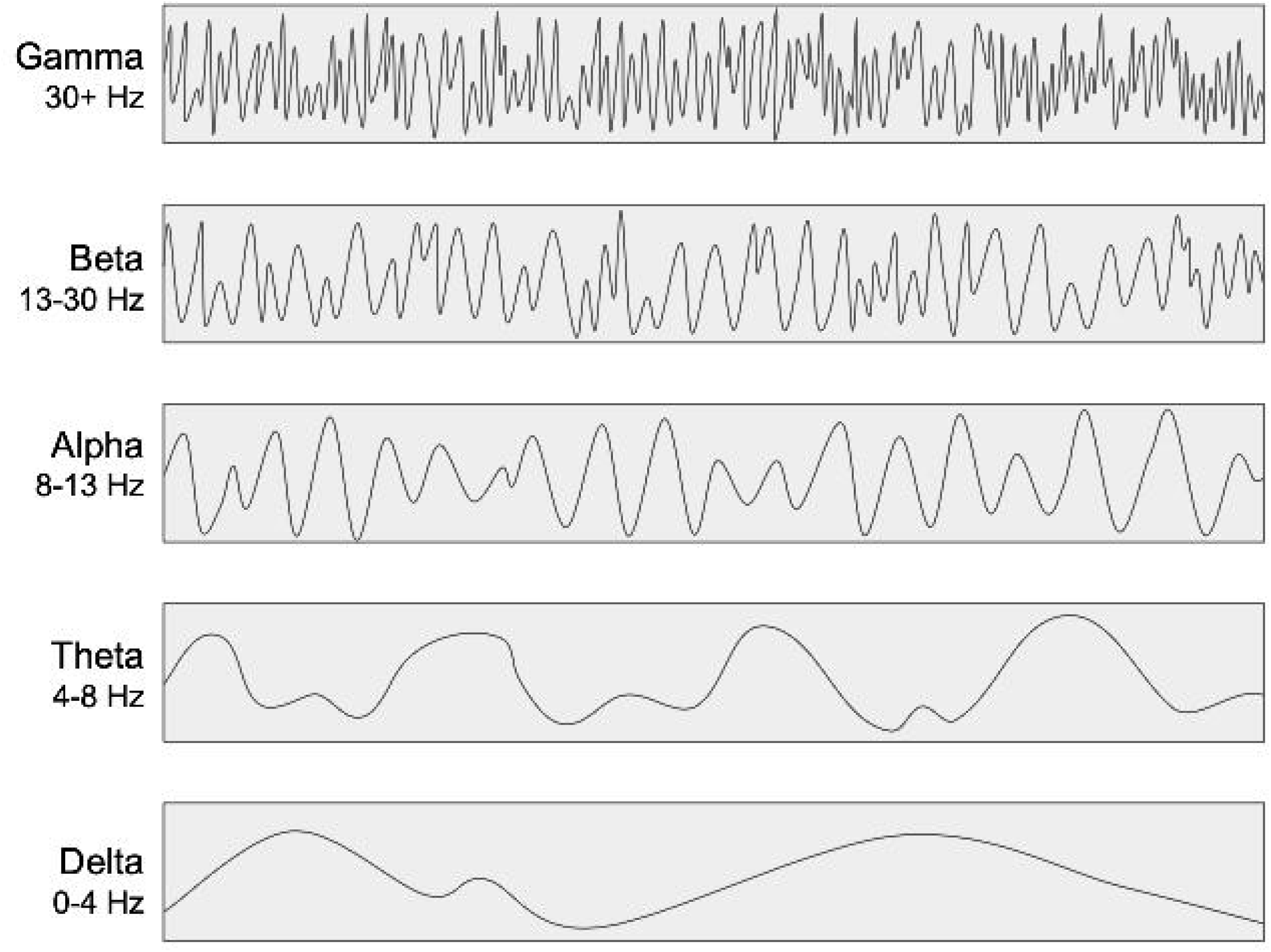
EEG Frequency Bands. This visualization demonstrates the differing frequencies of the various EEG bands. Created in google slides.

Neurofeedback originated in the 1960s for EEG (Kamiya, 2011), and early 2000s for fMRI (e.g., DeCharms et al., 2004; Yoo & Jolesz, 2002). Similar to biofeedback, it consists of relaying brain data (i.e., the target measure) to a participant while they perform a task. The participant may be given instructions to modulate the target by any number of strategies, or they may be given a specific strategy and told that correct application will be indicated by changes in their brain data. This neurofeedback condition may be compared to control conditions, wherein participants are presented with data from other brain regions not affected by the strategy (‘alternative ROI control’), or from other participants (‘yoked’ sham). Given well-designed controls (Sorger et al., 2019), neurofeedback can provide more substantive evidence that a brain region or network is involved in a process (for a complete account, neurostimulation methods may be most optimal). In addition to providing mechanistic insights into mental processes (Kvamme et al., 2022), neurofeedback allows participants to manipulate those processes. Neurofeedback has been used in many different applications, from the regulation of chronic pain (deCharms et al., 2005), to attentional training (typically involving prefrontal regions) (DeBettencourt et al., 2015; Wang & Hsieh, 2013), to stress reduction (typically involving the amygdala) (Hellrung et al., 2018; Nicholson et al., 2017; Young et al., 2017). It is often considered to ‘enhance’ learning, leading to improved outcomes (Haugg et al., 2021; Kadosh & Staunton, 2019). Researchers often conduct a single session of neurofeedback and then evaluate behavioral outcomes days, weeks or months later ((Pamplona et al., 2023; Ros et al., 2013); though, several studies have leveraged repeated sessions (Dekker et al., 2014; Mehler et al., 2018). Overall, there is promising evidence for the clinical mental health benefits of EEG and fMRI neurofeedback (Roy et al., 2020; Trambaiolli et al., 2021; Van Doren et al., 2019; c.f. Thibault et al., 2018). Thus researchers have proposed that the clinical benefits of mindfulness (as well as cognitive benefits) could be enhanced or facilitated by neurofeedback (Brandmeyer & Delorme, 2013; Brandmeyer & Reggente, 2023).

Starting in the early 2010s, neurofeedback concurrent with mindfulness meditation has been gaining popularity, and it is often referred to as mindfulness-based neurofeedback (mbNF). The purpose of this paper is to systematically review the literature and thus answer two main questions. First, can participants learn to modulate brain targets through mindfulness meditation practice, providing evidence of their involvement in meditation? Second, what are the behavioral and brain outcomes of mbNF? By reviewing the literature, there also are opportunities to discuss methodological limitations. The CRED-NF checklist (Ros et al., 2020), could be a crucial initial step towards standardizing current methodological and outcome reporting practices. The CRED-NF checklist includes preregistration, sample size justification, control group, double-blinding, whether or not participants used a strategy, artifact removal, feedback specification, regulation success (target engagement), brain and behavioral outcomes, and more. We evaluate the quality of studies herein based on the CRED-NF checklist. The present review only examines controlled lab-based EEG and fMRI studies (consumer-grade EEG studies are not reviewed, see Methods).

## 2 Methods

PRISMA guidelines were followed in this review (**Supplement 2**) (Page et al., 2021).

### 2.1 Inclusion and Exclusion Criteria

Studies that employed EEG or fMRI neurofeedback concurrently with mindfulness meditation were included. Specifically, studies were selected that claimed to employ mindfulness meditation, and we then evaluated whether the meditation met our definition of mindfulness. For the purposes of this review, mindfulness meditation is defined as meditation practice with the aim of cultivating non-judgmental attention to the present moment, including both focused attention (FA) and open monitoring (OM) practices (Lutz et al., 2008). FA and OM are distinct practices, but both are taught in mindfulness interventions like MBSR (Santorelli, 2014) and involve purposeful redirection of attention to the present moment (Britton et al., 2019; Dahl et al., 2015). Other meditation (e.g. transcendental, compassion) was not included. Exclusion criteria included lack of EEG or fMRI neurofeedback, lack of mindfulness meditation, lack of concurrent neurofeedback and mindfulness meditation, and non-empirical status (e.g., reviews). Studies with consumer-grade EEG devices were considered beyond the scope of this review, as there were substantial such studies (33), and consumer-grade devices don’t allow sufficient insight into brain mechanisms. No relevant conference papers or dissertations were identified.

### 2.2 Systematic Search

A search of PubMed, Web of Science, PsycInfo, and Scopus, was completed on November 11, 2023. Databases were identified based on previous mindfulness systematic reviews and meta-analyses (Goldberg et al., 2018; Treves et al., 2019). Search terms were “(mindfulness OR meditation) AND (neurofeedback OR neural feedback OR neuro feedback)”. We additionally searched reference sections of included papers.

### 2.3 Study Selection

All studies were first screened for duplicate publications. Next, all abstracts were screened, including studies based on two main criteria: full report of an empirical study (examples of excluded articles were review papers, protocol papers, book chapters, and conference proceedings) and content relevance (based on above stated inclusion/exclusion criteria). Then remaining studies were screened by reviewing the methods section and full paper to further evaluate the presence of inclusion criteria. Determination of inclusion was established in cases of disagreement by consulting with the first author.

### 2.4 Coding

Records were grouped according to neuroimaging technique (i.e., EEG or fMRI). Two reviewers (KDG & EW) independently evaluated each EEG study and its characteristics, and two reviewers (INT & ZB) independently evaluated each fMRI study and its characteristics. The studies were coded for sample, targets, neurofeedback details, control conditions, target engagement, neural outcomes and behavioral outcomes (**Tables 1** and **2**).

Target engagement was defined as ‘whether or not the neurofeedback target was modulated’, whereas neural outcomes are changes in other neural measures not targeted in the study neurofeedback protocol. Behavioral outcomes may consist of outcomes like state mindfulness reported after the scan, or more distal but related outcomes (e.g. cognitive performance on a separate task).

### 2.5 Bias and quality coding

No automation tools were used. Papers were coded independently to limit reviewer bias. Risk of bias in the studies was not quantified given the limited number of RCTs. Instead, we coded studies based on the CRED-NF checklist (Ros et al., 2020), reporting whether recommended items were present in the studies (**Table S1**).

## 3 Results

### 3.1 Search Results

A Prisma flow diagram is shown in **Figure 3**. The search yielded 676 records across four databases. After removing duplicates and excluding based on title and abstract, full texts were reviewed for the remaining 114 studies. The final sample included 19 studies with 15 independent samples representing 419 participants.

**Figure 3:**
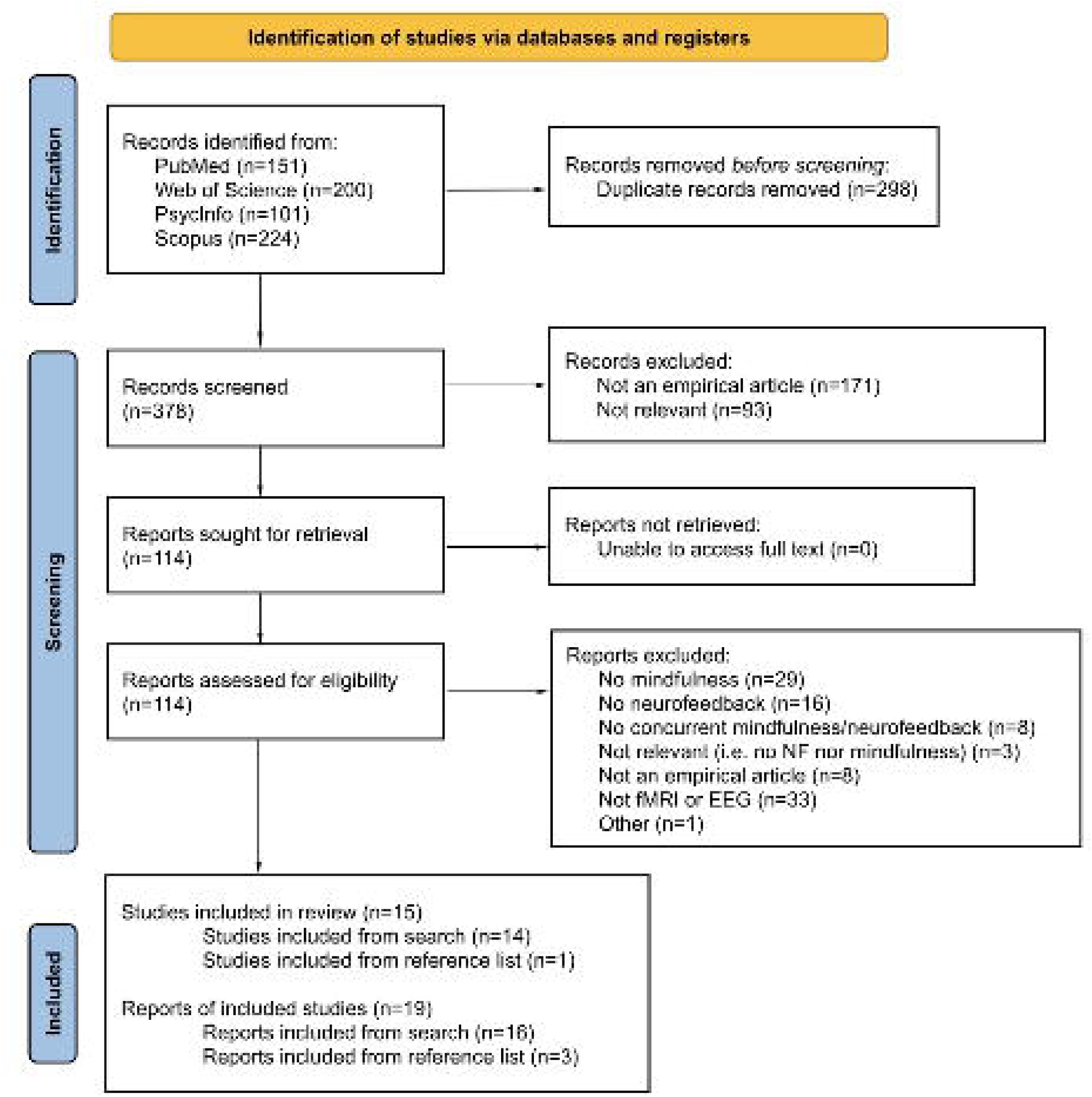
PRISMA flow diagram depicting number of identified and evaluated articles for concurrent mindfulness and fMRI or EEG neurofeedback procedures. *Note.* Studies refer to unique samples, while reports refer to publications on said samples. Our review identified four samples which corresponded to more than one published report, as indicated in this flowchart and in the study summary tables (Tables 1 and 2).

**Table 1.**
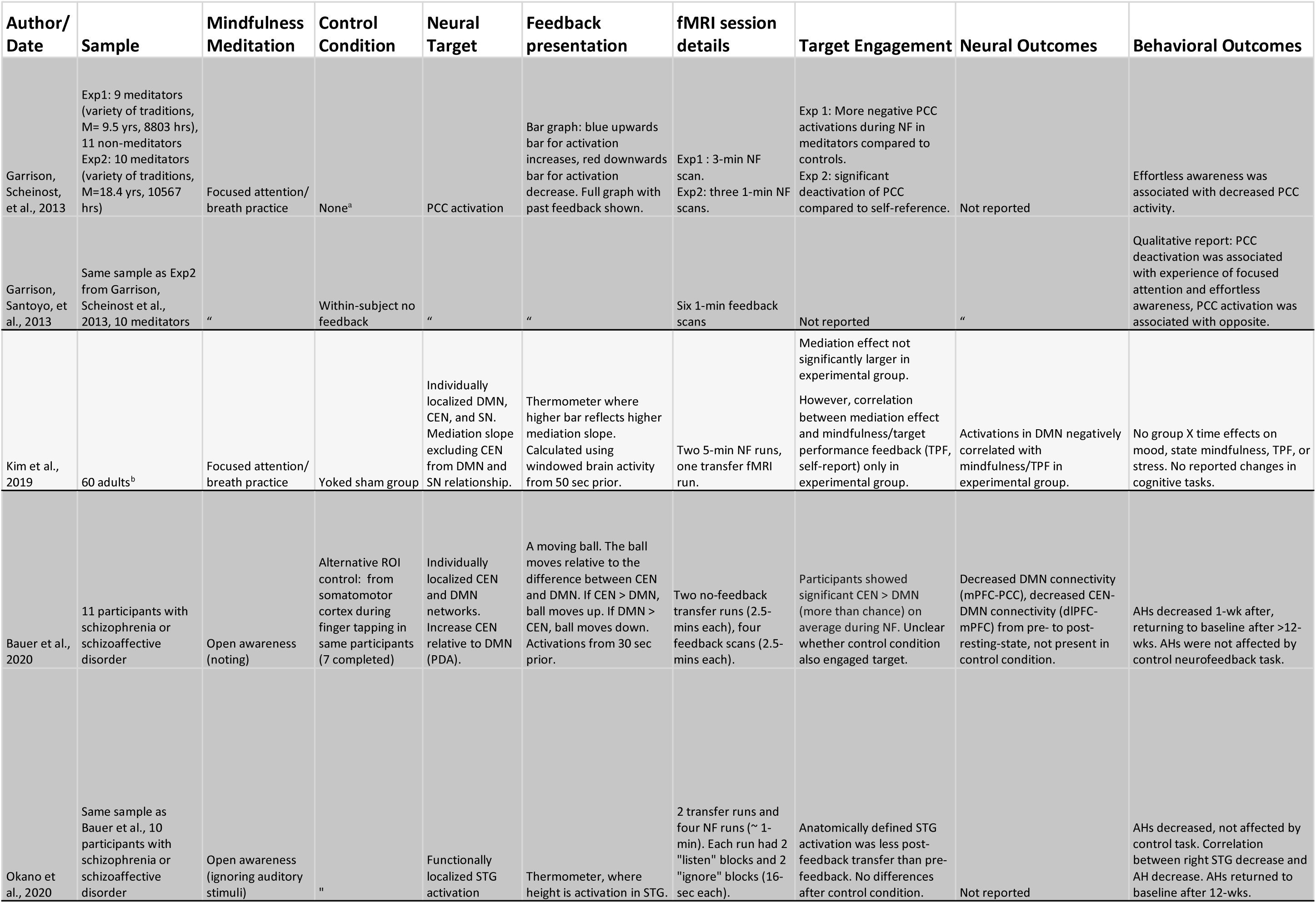
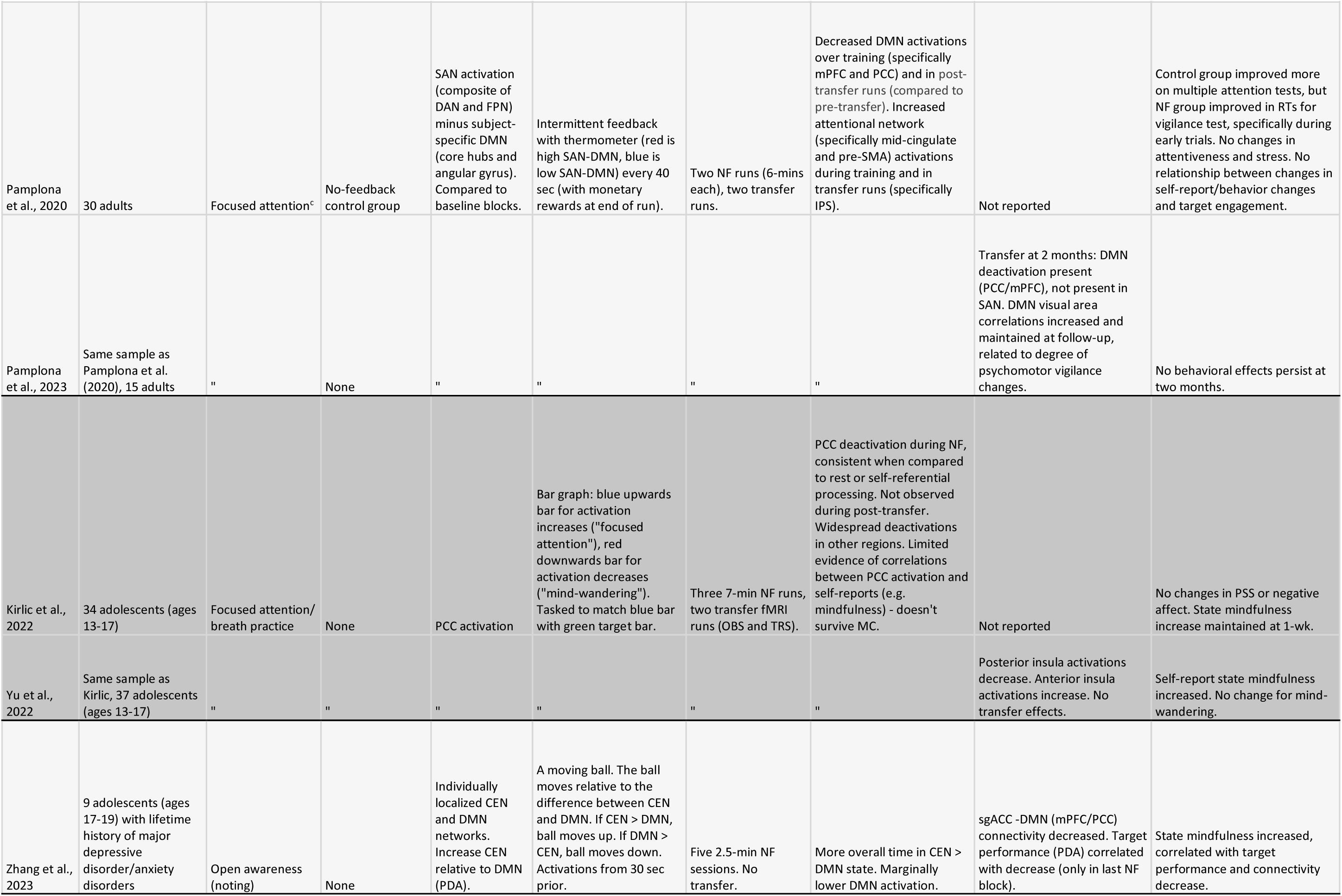
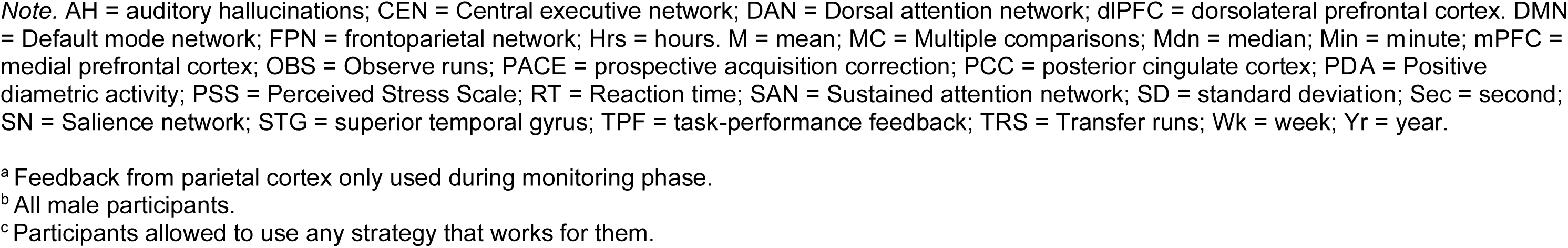
Summary of studies of fMRI-based neurofeedback with concurrent mindfulness practice.

**Table 2.**
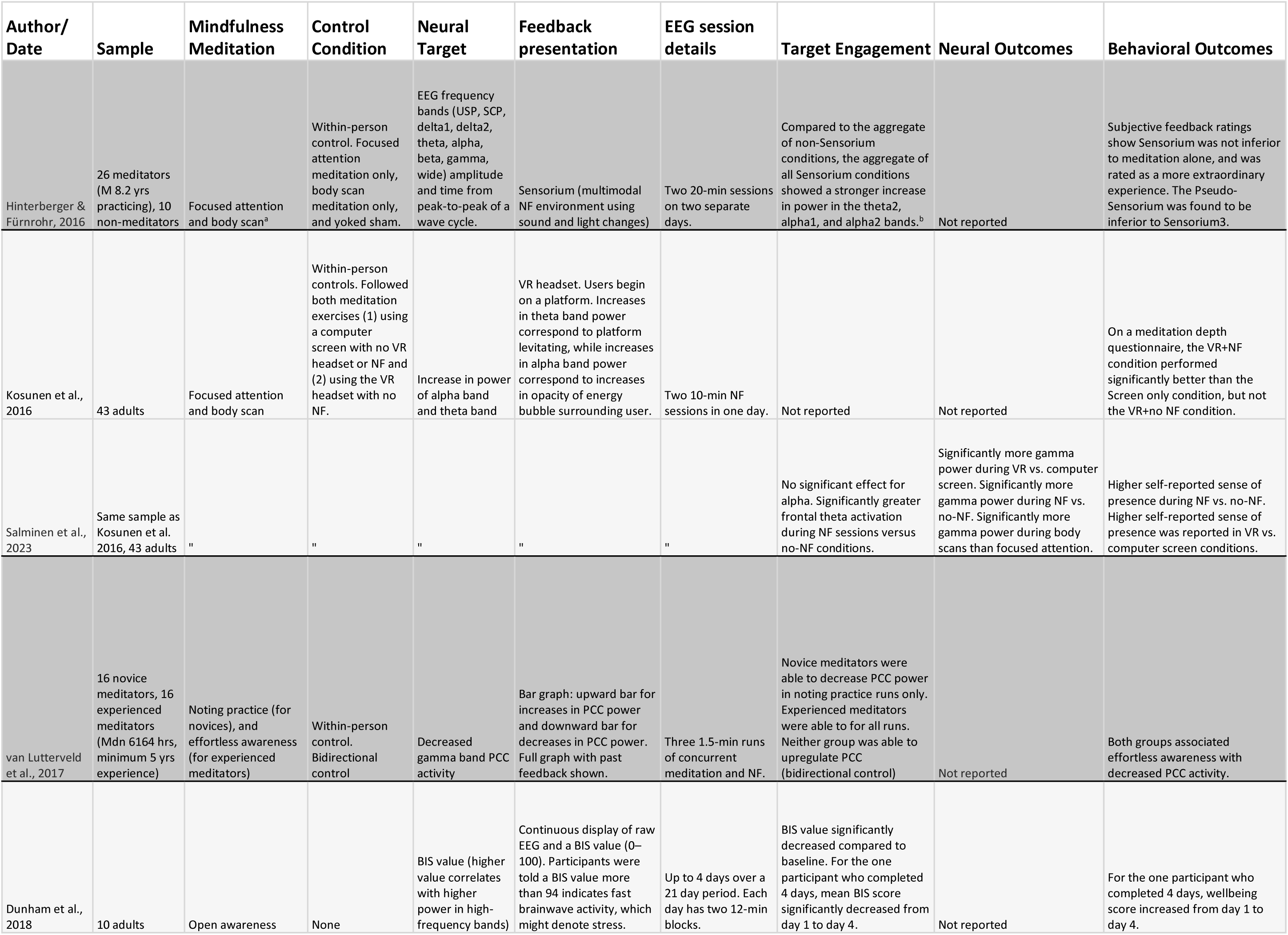
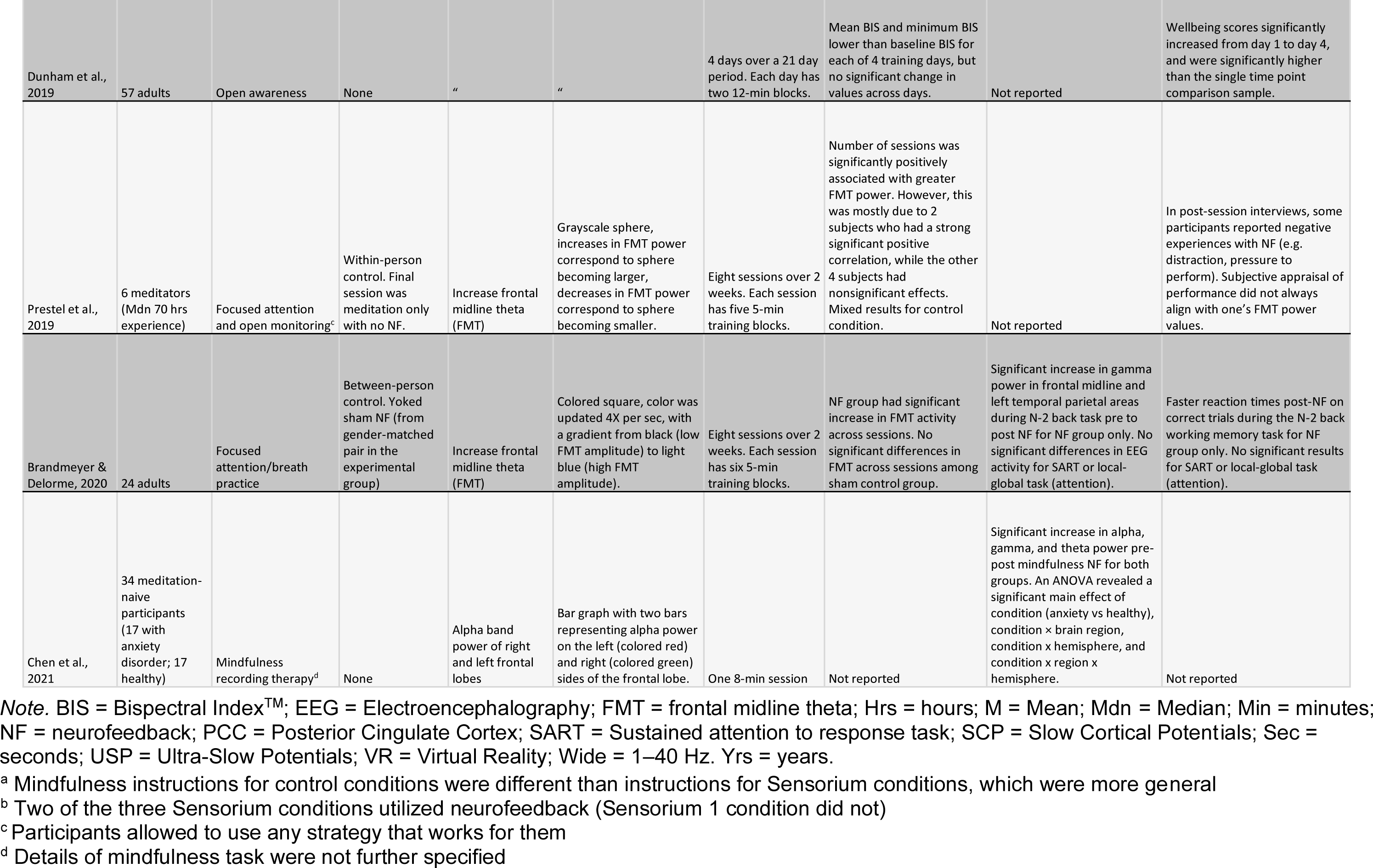
Summary of studies of EEG-based neurofeedback with concurrent mindfulness practice.

### 3.2 fMRI Studies

#### 3.2.1 Summary

We identified 10 reports of fMRI neurofeedback including 7 unique samples (**Table 1**). There were two studies that used different neurofeedback protocols with the same sample, patients with schizoaffective disorders (Bauer et al., 2020; Okano et al., 2020). In total, there were 177 unique participants, and most samples were at or below 30 participants (**Figure S1 & S2**). Studies employed focused attention and open monitoring meditations. The first study published was Garrison et al. (2013a), a breath-focused attention study, which was also the only study to involve experienced meditators. Garrison et al. (2013b) examined qualitative reports from meditators who were asked to explore correspondence between brain signals and meditation experiences. A typical open monitoring protocol asked participants to label thoughts and feelings as they came up (Bauer et al., 2020). Only one study reported participants’ actual mental strategies during neurofeedback regulation (Pamplona et al., 2020), and found a variety of strategies employed, some of which weren’t typical mindfulness. Control conditions encapsulated yoked sham, alternate ROI, and mindfulness meditation without feedback, but many studies did not include controls (**Figure S3 & S4)**. Only two studies examined between-subjects controls (**Figure S5**) (Kim et al., 2019; Pamplona et al., 2020), and transfer runs (**Box 1**) were inconsistently used across studies. Zhang et al. (2023) was the only study to examine adolescents. Qualitative assessments of the mbNF experience are found in **Table S3.**

##### Box 1: Neurofeedback Terms

###### Bidirectional control

Testing whether participants may modulate a neurofeedback target in both directions. For example, decreasing the DMN by meditating, and increasing the DMN by ruminating.

###### Calibration

A preceding block of non-neurofeedback data used for the neurofeedback target estimates, typically eyes-open rest.

###### Control, Alternate ROI

Typically, feedback is given from a region or network that is not related to the task. <u>Control, Yoked Sham:</u> Feedback is presented to a control participant from an experimental participant. This feedback is controlled for in terms of perceived reward but not contingent on a control participant’s performance.

###### Functional/individual localization

Determining a brain area or network based on data from the participant. An example is conducting resting-state fMRI before the neurofeedback task, which can be used to extract intrinsic networks that are correlated at rest.

###### Intermittent vs continuous

Intermittent, or delayed, feedback is feedback presented after regular intervals, not concurrently with task. Continuous, or real-time, feedback is feedback presented throughout the task (e.g. every second). May involve different attentional demands (Hellrung et al., 2018).

###### Offline artifact correction

Estimates of motion or physiology are corrected for or tested for in post-processing. <u>Online artifact correction:</u> Estimates of motion or physiology are included in real-time models (e.g. GLMs), so feedback is not presented based on those artifacts.

###### Target

The brain measure relayed to participants.

###### Target engagement

A test of whether participants successfully learned to modulate the target brain measure, may consist of examining overall levels of target, change in target, or target performance in transfer runs.

###### Transfer run

A neuroimaging run where participants perform the neurofeedback task without any feedback presented. Transfer tasks after feedback can be used to assess whether learning has occurred.

#### 3.2.2 fMRI Targets

The majority of studies employed activation-based, default-mode network targets (**Figure 4**). There was some variety in the specification of the DMN. Two studies targeted PCC activity (Garrison et al., 2013; Kirlic et al., 2022, Yu et al., 2022). Multiple studies used individualized networks generated from independent component analysis of resting-state scans (Bauer et al., 2020; Kim et al., 2019; Pamplona et al., 2020, 2023). These studies combined network activations from not only the DMN but also attentional networks like the dorsal attention network (Pamplona et al., 2020, 2023), the salience network (Kim et al., 2019), and the CEN (Bauer et al., 2020). One study used a task to functionally localize the superior temporal gyrus (STG) in order to modulate auditory processing (Okano et al., 2021). Kim et al. (2019) was the only study which used a connectivity-like target, and they examined the direct effect from DMN-SN, excluding the influence of the CEN (as the CEN was proposed to be involved with the visual feedback monitoring and not the meditation). Target measures did not appear to depend on whether participants performed focused attention vs open monitoring.

**Figure 4:**
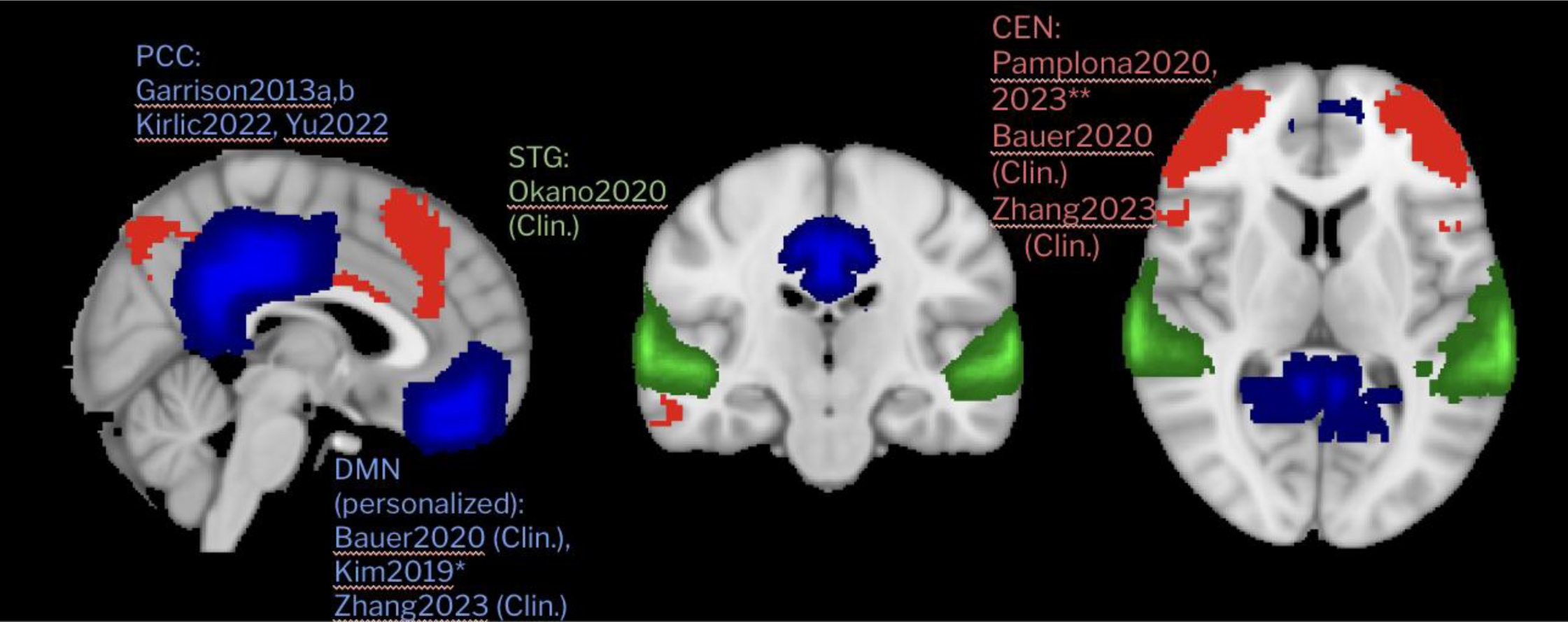
fMRI targets for neurofeedback. In blue, default-mode network (DMN) regions. In green, the superior temporal gyrus (STG). In red, the central executive network (CEN). Networks are taken from the Yeo atlas (Yeo et al., 2011), and STG was extracted from the Harvard-Oxford cortical atlas. *Kim used DMN-SN slope **Pamplona used sustained attention network.

There were different approaches to computing the target measure. Multiple studies used baseline periods in the same fMRI scan to scale and baseline the neurofeedback target measure (e.g., 30 seconds of rest; Bauer et al., 2020; Okano et al., 2020; Zhang et al., 2023). Garrison et al., 2013a, 2013b used a self-reference task. Some studies used online motion artifact correction, and two studies additionally conducted online correction for physiological signals like breathing and heart rate (Kirlic et al., 2022; Yu et al., 2022). No differences were observed between neurofeedback and rest in terms of motion or physiological signals.

Feedback was displayed visually to participants during fMRI in all cases. Some studies used delayed feedback, e.g. continuous feedback but with a time lag (Bauer et al., 2020; Kim et al., 2019), and one used intermittent feedback (Pamplona et al., 2020). One study displayed a continuously updating graph containing past values of the measure (Garrison et al., 2013). Negative and positive feedback was shown. One study incorporated rewards for target engagement (Pamplona et al., 2020).

#### 3.2.3 fMRI Target Engagement

All studies evaluated target engagement. Garrison et al. (2013) found that meditators showed more negative PCC (the target) activation than controls. Two studies examined the amount of time spent in a ‘correct’ brain state (CEN > DMN), and found above chance engagement for the group (Bauer et al., 2020; Zhang et al., 2023). Another study examined change over neurofeedback blocks in activation, and found significant decreases in the DMN (Pamplona et al., 2020), but no increases in their individually defined attentional networks. Transfer runs can also be used to assess whether target engagement or learning has taken place. Kirlic et al. (2022) observed decreased PCC activation during neurofeedback compared to control tasks, but not in the post-transfer run. Although there seems to be consistent evidence of DMN deactivation during neurofeedback, the only study to use a sham control and a large (n∼60) sample did not find evidence for target engagement (mediation slope between the DMN-SN) (Kim et al., 2019). This and the transfer results from Kirlic and colleagues make it unclear whether participants have learned to modulate their brain networks.

#### 3.2.4 Neural Outcomes

Many studies examined the possibility of neural changes due to neurofeedback. Some reports focused on DMN-based connectivity assessed during resting-state fMRI before and after neurofeedback. There are indications of reduced within-DMN connectivity (e.g. between the MPFC and PCC), as well as more negative correlations between DMN and CEN (Bauer et al., 2020). Other studies found reduced DMN connectivity with the sgACC (sometimes considered part of the DMN; Zhang et al., 2023), reduced DMN connectivity with the STG (Okano et al., 2020), and increased DMN-visual area connectivity (Pamplona et al., 2023) even at a 2-month follow-up assessment. Of the findings, only the DMN-STG finding was established as specific to the mindfulness-based neurofeedback, as Okano and colleagues did not find DMN-STG changes in an alternative ROI control session (finger tapping). However, Zhang et al. (2023) found associations between DMN-sgACC connectivity decreases and target engagement and state mindfulness in their small sample.

Researchers also examined activations in non-target and target brain regions. Yu et al. (2022) found that neurofeedback increased anterior insula activations, and decreased posterior insula activations, without any transfer effects. This could reflect changes in interoceptive processing during mbNF. Kim et al. (2019) found that activations in the DMN negatively correlated with state mindfulness, but only in the experimental group and not the sham feedback group.

#### 3.2.5 State Mindfulness

To assess whether participants are learning from neurofeedback, studies also tested whether they experienced increases in state (or momentary) mindfulness, as reported after the scans. State mindfulness assessments typically involved questions about present-focused awareness of the mind and body (Tanay & Bernstein, 2013). Kim et al. (2019) found no mbNF vs control group effects on state mindfulness or self-report target efficacy. However, two uncontrolled studies found increases in state mindfulness after neurofeedback (Kirlic et al., 2022; Zhang et al., 2023).

#### 3.2.6 Behavioral Outcomes

A central motivation for many of the studies was the possibility of beneficial behavioral or self-reported outcomes. Pamplona et al., (2020) observed improvements in reaction time on a vigilance test, but this was not maintained at the 2-month follow-up (Pamplona et al., 2023). They also did not observe a correlation between target engagement and vigilance reaction time. Kirlic et al. (2022) did not observe any changes in perceived stress or negative affect after neurofeedback. Bauer et al. and Okano et al. observed decreases in auditory hallucinations that were not present after a control neurofeedback task, however, the protocols were conducted on the same sample and the same baseline was used for both. Overall, there is limited evidence for mindfulness-based fMRI neurofeedback benefits as yet.

#### 3.2.7 Clinical Applications

Three studies conducted neurofeedback with small clinical samples (Bauer et al., 2020; Okano et al., 2020; Zhang et al., 2023). Bauer and Okano et al. examined neurofeedback in the same 10 individuals with schizophrenia, and found decreases in auditory hallucinations - although these changes were not sustained at 12 weeks. Zhang et al. examined neurofeedback in 9 adolescents with affective disorder history, and found decreases in sgACC-DMN connectivity which is heavily implicated in adolescent depression (Chai et al., 2016); though symptom changes were not assessed. These studies can be considered pilots–focused mostly on establishing feasibility of the neurofeedback protocols in clinical samples.

#### 3.2.8 Quality (CRED-NF)

In general, the control conditions in fMRI studies (within and across-subjects) were lacking, with only one study involving adequate controls and reporting. Reporting of feedback specifications, target engagement (in the feedback condition), data processing methods, etc. was present across the vast majority of studies. Few studies conducted preregistration, power analyses, or made their data/code open access. A full table may be found in **Table S2.**

### 3.3 EEG studies

#### 3.3.1 Summary

Nine reports of EEG neurofeedback during mindfulness meditation were identified, corresponding to eight unique samples (**Figure S1 & S2**). In total, there were 242 unique participants, and all samples were adults. Multiple samples looked at meditators, or compared meditators to non-meditators (Hinterberger & Fürnrohr, 2016; Prestel et al., 2019; van Lutterveld et al., 2017). Only one sample included a clinical population (anxiety disorders; Chen et al., 2021) and only two samples did not include a control condition (Chen et al., 2021; Dunham et al., 2018; Dunham et al., 2019). All control conditions were within-subject with the exception of Brandmeyer and Delorme (2020) (**Figure S5)**. However, the control conditions varied; most were compared to some form of meditation without neurofeedback and others included yoked shams (**Figure S3 & S4**) (Hinterberger & Fürnrohr, 2016; Brandmeyer & Delorme, 2020). The most common types of meditation were focused attention, body scan, and open monitoring. The terminology for the type of meditation was not always consistent, and we used specific reporting from studies to classify meditation types. That said, reporting on specific mindfulness instructions was not always clear (Chen et al., 2021) and participants were sometimes allowed to use various strategies (Prestel et al., 2019). It is also important to note that even within a single study, the instructions of the control condition mindfulness did not always match the instructions of the active NF session (Hinterberger & Fürnrohr, 2016). Qualitative assessments of the mbNF experience are found in **Supplement Table 2.**

#### 3.3.2 EEG Targets

Almost all studies used changes in frequency band power as their neural target (**Figure 5**); the most common was alpha and theta, though some studies used gamma (van Lutterveld et al., 2017) or Bispectral Index^TM^ (BIS) value, which is an EEG technique most commonly used to measure depth of consciousness for patients under general anesthesia (Dunham et al., 2018; Dunham et al., 2019). Multiple studies focused on more than one frequency band (Hinterberger & Fürnrohr, 2016; Kosunen et al., 2016; Salminen et al., 2023). Some studies focused on whole brain frequency band power, while others looked at frontal midline sites (Brandmeyer & Delorme, 2020; Prestel et al., 2019) or source localized areas like the PCC (van Lutterveld et al., 2017). The density of EEG ranged from high density 128-channel (van Lutterveld et al., 2017) to extremely low density Bispectral Index, which generally has 2-4 channels though the exact number of channels was not reported in this case (Dunham et al., 2018; Dunham et al., 2019). Notably, the way the target was calculated varied, even within a sample. For example, Salminen et al. (2023) calculated theta power from an average of two electrodes (F3 and F4) and alpha power from an average of all electrodes (F3, F4, C3, C4, P3, and P4). Other studies used an independent component analysis (ICA) to calculate the target (Chen et al., 2021; Prestel et al., 2019). Interestingly, the only two samples that had the same neural target (frontal midline theta) calculated the feedback differently, with Brandmeyer & Delorme (2020) using the signal from a single frontal electrode (Fz) while Prestel et al. (2019) used an ICA to determine frontal midline theta. Almost all feedback was displayed visually, most commonly on some sort of screen, though virtual reality was also used (Kosunen et al., 2016; Salminen et al., 2023). One study used both sounds and light changes as their feedback presentation (Hinterberger & Fürnrohr, 2016). Positive and negative feedback was shown for all studies.

**Figure 5.**
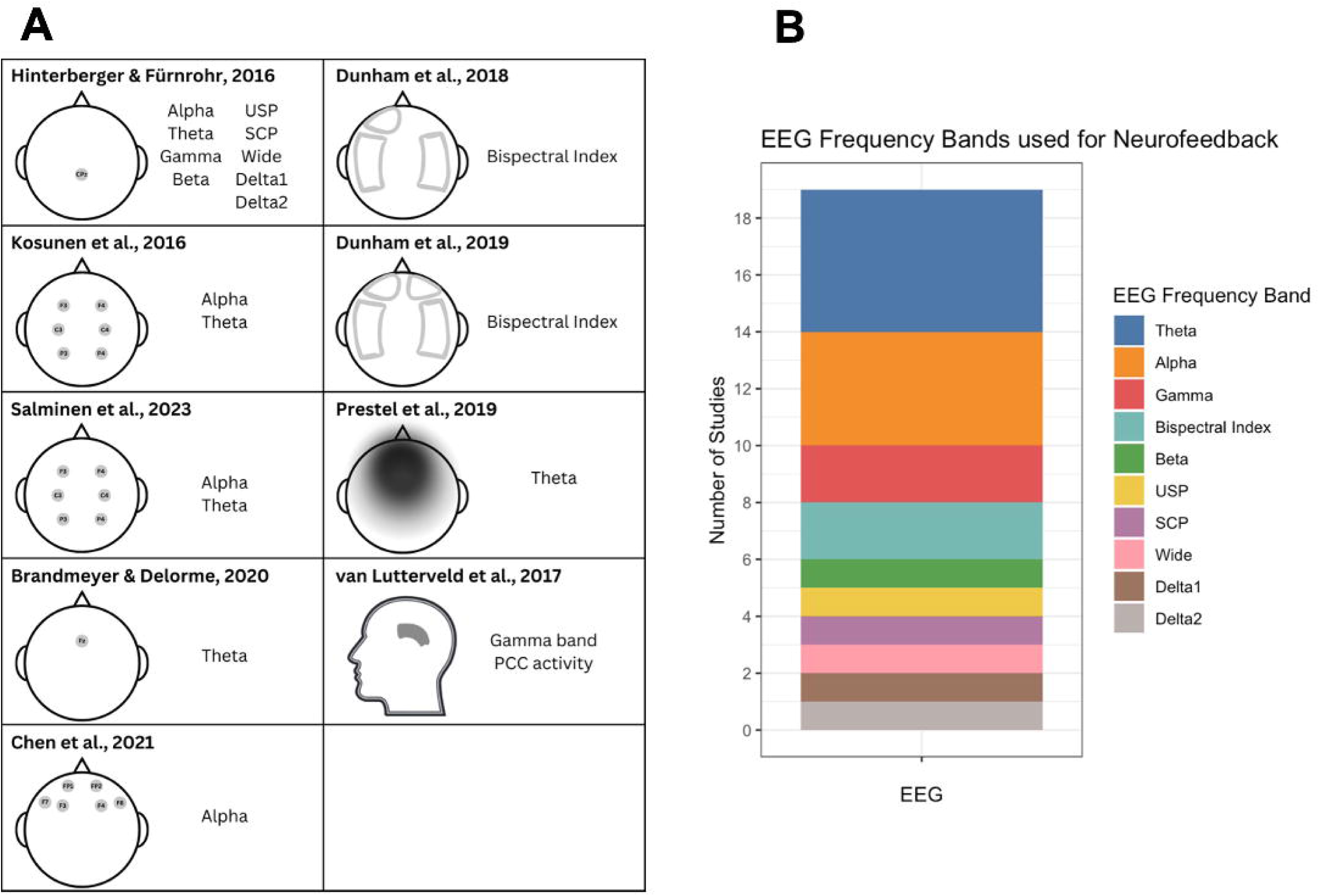
This figure displays the variety in methods of calculating EEG-neurofeedback. In **A),** the spatial layouts of the neurofeedback targets are displayed. Hinterberger & Fürnrohr, 2016 calculated alpha, theta, beta, gamma, USP, SCP, wide, delta1, and delta2 from CPz. Kosunen et al., 2016 calculated theta and alpha using an average from F3, F4, C3, C4, P3, and P4. Salminen et al., 2023 calculated theta power using an average F3 and F4, while alpha power was calculated as an average of F3, F4, C3, C4, P3, and P4. Brandmeyer & Delorme, 2020 used Fz to calculate frontal midline theta. Chen et al., 2021 used FP1, F3, F7 and FP2, F4, F8 to calculate alpha of the left and right frontal lobes, respectively. Dunham et al., 2018 placed the BIS sensor on the left forehead and the temporal fossa. Dunham et al., 2019 placed the BIS sensor on the left or right forehead and the temporal fossa. Prestel et al., 2019 used independent component analysis (ICA) with a 32-channel setup to calculate frontal midline theta. van Lutterveld et al., 2017 used source estimation with a 128-channel setup to determine Gamma band PCC activity. In **B),** the frequency bands used by each study are displayed. Theta was used the most by five studies, closely followed by alpha, which was used in four studies.

#### 3.3.3 EEG Target Engagement

Not all papers reported on target engagement (Kosunen et al., 2016; Chen et al., 2021). For those that did, the way the target engagement was reported differed. For example, some studies looked at activation compared to baseline (Dunham et al., 2018; Dunham et al., 2019; van Lutterveld et al., 2017), compared to no NF conditions (Salminen et al., 2023), or across sessions using different linear models (Prestel et al., 2019; Brandmeyer & Delorme, 2020). The difference in reporting and measures used makes synthesizing results difficult, as well as the fact that support for target engagement was mixed. The strongest support was for an impact on theta (Brandmeyer & Delorme, 2020; Hinterberger & Fürnrohr, 2016; Salminen et al., 2023; Prestel et al., 2019), while alpha had both significant and null results (Hinterberger & Fürnrohr, 2016; Salminen et al., 2023). However, it is important to note Hinterberger and Fürnrohr’s (2016) significant results were collapsed across multiple experimental conditions, one of which did not include NF. Others found significant changes in BIS values compared to baseline (Dunham et al., 2018; Dunham et al., 2019), but the significant decrease from day one to day four (Dunham et al., 2018) did not hold up in a larger sample (Dunham et al., 2019). For those who did find significant results across days, the significant effect was in one case driven by two participants (Prestel et al., 2019), and was stronger when certain “non-responders” were excluded (Brandmeyer & Delorme, 2020). It is also important to note the spatial limitations of neural targets using EEG, given that spatial resolution of EEG is limited even when using source localization. For example, the one study that used source localization of the PCC found that over 80% of runs examined had significant correlations between the right lateral occipital cortex and the PCC, though less than 36% of runs showed significant correlation between the left supplementary motor area and the PCC (van Lutterveld et al., 2017). Correlations between 40-57Hz PCC time series and delta, theta, alpha, and beta were calculated, but never surpassed more than 40% of runs showing significant correlations. The frequency specific to the PCC may be more accurate than the source localization, which may be capturing signals from occipital regions of the brain more broadly.

#### 3.3.4 Neural Outcomes

Only four studies out of nine reported on other neural outcomes. All four studies examined EEG bands beyond the bands of the neural target, focusing on: (a) other frequency bands during NF and/or control (Salminen et al., 2023; van Lutterveld et al., 2017), (b) frequency bands at rest before and after NF (Chen et al., 2021), and (c) frequency bands in tasks before and after NF (Brandmeyer & Delorme, 2020). The following results discuss frequency bands that were not the target of NF. There is support that alpha, theta, and gamma power significantly increase from pre to post mindfulness NF (Chen et al. 2021), as well as support for gamma increases during NF compared to no-NF (Salminen et al., 2023). The delivery of NF (Virtual Reality (VR) > computer screen) and the type of meditation (body scan > focused attention) can also have an impact on the level of gamma (Salminen et al., 2023). There is mixed support for NF’s effect on cognitive tasks, with no significant changes found for attentional tasks, but a significant increase in gamma power during a working memory task done before and after NF for those that received NF (Brandmeyer & Delorme, 2020).

#### 3.3.5 Behavioral Outcomes

All but one study reported on some sort of behavioral outcome. Most focused on how individuals felt doing the mbNF (Hinterberger & Fürnrohr, 2016; Kosunen et al., 2016; Prestel et al., 2019; Salminen et al., 2023), while others looked at changes in well-being (Dunham et al., 2018; Dunham et al., 2019) or even performance on cognitive and attentional tasks (Brandmeyer & Delorme, 2020). When looking at immersive ways to deliver neurofeedback, such as the Sensorium or VR, findings suggest participants find these types of modalities more extraordinary, more engaging, have more positive experiences and less negative experiences compared to audio/visual guided meditations without VR or Sensorium enhancements (Hinterberger & Fürnrohr, 2016; Kosunen et al., 2016; Salminen et al., 2023). However, this difference is not always due to the addition of neurofeedback, as control conditions with these enhancements with yoked sham or no NF did not always show significant differences (Hinterberger & Fürnrohr, 2016; Kosunen et al., 2016). Accordingly, some participants report that too much focus on the NF can be distracting and lead to poorer performance; however, it is interesting to note that subjective experience of performance did not always align with objective performance, as measured by frontal midline theta (Prestel et al., 2019). Beyond experiences during mbNF, studies have found that wellbeing scores increase from baseline to completion of all sessions, and are significantly higher at completion than a control group who received no mbNF (Dunham et al., 2018; Dunham et al., 2019). There is also some evidence to suggest NF may help improve performance on memory tasks (NF group compared to sham control had faster reaction times post-NF for correct trials during the N-2 back working memory task) (Brandmeyer & Delorme, 2020). However no significant effects were found for attention tasks.

#### 3.3.6 Clinical Applications

Only one study examined EEG neurofeedback in a clinical sample of 17 individuals with anxiety disorders (Chen et al., 2021). Compared to healthy controls, anxious subjects exhibited initial lower power in alpha, theta, and gamma. After NF, anxious subjects significantly increased power across all bands in all brain areas. Chen et al. (2021) suggest that the increase in gamma power indicated a reduction in anxiety symptoms, though they did not report on changes in subjective measures of anxiety. ANOVAs revealed interactions of condition vs brain region/hemisphere, but the direction of differences were not reported. Although the remaining studies reported on non-clinical samples, Dunham et al., 2018 and Dunham et al., 2019 examined well-being as a target for mindfulness neurofeedback among physicians/nurses, a group within which stress and emotional exhaustion are common (Dunham et al., 2018). Participants’ subjective well-being was found to improve following the mbNF, suggesting that even in non-clinical samples, NF may be a promising avenue to increase well-being.

#### 3.2.8 Quality (CRED-NF)

There were control conditions in the majority of the EEG studies, but they typically lacked blinding (sham). Reporting of feedback specifications and target engagement was common. Few studies reported artifact correction. Few studies conducted preregistration, justified their sample sizes or made their data/code open access. A full table may be found in **Table S3.**

## 4. Discussion

Mindfulness meditation consists of purposefully bringing one’s attention back to the present moment, and cultivating an open-minded and non-judgmental attitude (Creswell et al., 2017). Though mindfulness meditation is increasingly used for promoting mental health, there are many open questions about its neural bases. In this review, we investigate a promising tool for understanding the neural mechanisms of mindfulness, neurofeedback. Neurofeedback consists of relaying neural signals (the target) to the participant and examining if they can learn to modulate the signals (target engagement). Successful modulation provides evidence that a target brain region is involved in meditation. In addition, given the right targets, neurofeedback may help participants practice correctly and lead to better attention, deeper mindfulness, and positive behavioral outcomes. In this systematic review, we assess whether participants can modulate brain targets (insight into neural mechanisms) and whether participants benefit from the practice (behavioral outcomes). We included studies utilizing mindfulness meditation with concurrent EEG or fMRI feedback (i.e., mindfulness-based neurofeedback [mbNF]).

The search yielded 19 reports, with 15 independent samples. The earliest study was published in 2013, underscoring the nascency of the mbNF field (systematic inquiry of neurofeedback more generally extends back to the early 2000s for fMRIs and the 1960s for EEG). Studies used a wide range of targets across brain areas and frequency bands, and often reported different metrics of target engagement. Neurofeedback duration and number of runs varied (from single 15-min sessions to multiple weeks of training). Sample sizes were generally small, given the resource-intensive nature of neurofeedback. Few studies were RCTs, which are critical for establishing mbNF efficacy and testing mechanisms.

### 4.1 Brain Targets

One of the prominent neuroscientific theories of mindfulness posits that successful practice leads to downregulation of the DMN, perhaps most robustly the core hubs of the PCC and mPFC (Ganesan et al., 2022). The DMN has a well-established role in internally-generated, self-referential thought (Andrews-Hanna, 2012; Buckner et al., 2008). Mindfulness meditation involves recognizing self-referential thoughts, disengaging from them, and engaging in attention on an object like the breath. Thus, mindfulness may involve downregulating DMN activity. Accordingly, many fMRI studies of mbNF chose to target the DMN. Some studies calculated and displayed anatomically defined PCC activations compared to a control self-reference condition (Garrison, Santoyo, et al., 2013; Garrison, Scheinost, et al., 2013), whereas others used subject-specific, functionally derived maps of the DMN (Bauer et al., 2020; Kim et al., 2019; Okano et al., 2020; Pamplona et al., 2020; Zhang et al., 2023). Consistent downregulation of the DMN was found. Neurofeedback studies often included other networks like the central executive network (CEN) and salience network (SN). There is extensive reason to believe that DMN and other network interactions may be involved in mindfulness meditation, specifically in the switching between external and internal modes of attention (Hasenkamp et al., 2012; Mooneyham et al., 2016; Rahrig et al., 2022). Accordingly, studies relayed the participants’ difference between CEN and DMN (Bauer et al., 2020; Zhang et al., 2023), sustained attention networks and DMN (Pamplona et al., 2020, 2023), and the slope of the DMN-SN connectivity excluding the CEN (Kim et al., 2019). It is unclear when participants are given these multivariate measures which variable is being trained –one study found that the DMN was modulated and not the sustained attention network (Pamplona et al., 2020).

Researchers also examined neuroplastic changes dependent on their DMN-based neurofeedback, finding changes in DMN region connectivity with other brain areas like the anterior cingulate cortex (Zhang et al., 2023). These changes suggest that neurofeedback may modulate intrinsic features of the DMN, offering a key inroad to mitigate ruminative and depressogenic perseveration tendencies (Zhang et al., 2021).

Mindfulness meditation has been associated with power increases in alpha, theta and gamma waves during meditation (Chiesa & Serretti, 2010; Lee et al., 2018; Lomas et al., 2015; Stapleton et al., 2020). Alpha and theta power may correspond to shifting attention to internal sensations and thoughts (Lomas et al., 2015), whereas gamma power may reflect wider awareness (Lomas et al., 2015; Lutz et al., 2004; Stapleton et al., 2020). There is considerable evidence that gamma EEG activity can be contaminated by muscle activity (Muthukumaraswamy, 2013; Whitham et al., 2007). However, it is not necessary to disregard gamma power altogether, as long as multiple precautions are taken to remove muscle artifacts and confirm they are not correlated with data (e.g., van Lutterveld et al., 2017). Accordingly, the 9 EEG studies of mbNF selected alpha, theta and gamma targets. The most consistent evidence was for theta increases, specifically frontal midline theta, which is often an indicator of cognitive control (Cavanagh & Frank, 2014). Results were mixed when probing alpha and gamma power.

The study of EEG and fMRI have often been conducted in isolation, and each has advantages and disadvantages. EEG-neurofeedback is useful for precise temporal modulation as well as cost-effective application but lacks spatial specificity and may be susceptible to motor artifacts (Muthukumaraswamy, 2013; Whitham et al., 2007). fMRI-neurofeedback is useful for targeting specific brain regions with specificity, however, it is expensive and the underlying signals are slow to change. There have been meaningful efforts to develop EEG measures with spatial specificity. Frontal midline theta may be negatively correlated with DMN activation (Scheeringa et al., 2008; Prestel et al., 2018), while gamma may be positively correlated with DMN activation (Berkovich-Ohana et al., 2012). One neurofeedback study included, Van Lutterveld et al. (2017), specifically targeted activity in the PCC by using source localization. Yet, they did find that occipital cortex activity correlated heavily with localized PCC activity. It may be necessary to conduct EEG-fMRI fusion experiments to develop better measures. In a seminal paper, Keynan et al. (2019) created an EEG target measure of amygdala activity derived from machine learning based on simultaneous EEG-fMRI and showed that participants could modulate the target. The amygdala-EEG neurofeedback led to increases in emotional awareness and regulation and decreases in amygdala activation as measured by fMRI.

### 4.3 Brain Target Summary and Limitations

Extant research has, at times, corroborated neuroscientific theories of mindfulness; however, the majority of research did not include robust control conditions, which results in a lack of specificity. For example, decreases in DMN activation during neurofeedback does not indicate that participants are learning or that DMN deactivation is linked to mindful states. One possibility is that focusing on the display of the feedback itself may lead to DMN decreases. There is substantial evidence that engaging in external tasks leads to decreases in DMN activations (Raichle, 2015; Whitfield-Gabrieli & Ford, 2012), and likely changes in power as well (Fitzgibbon et al., 2004; Khader & Rösler, 2011). Another possibility is that mindfulness meditation leads to decreases in DMN activation, but that this process is implicit and beyond conscious control (thus, neurofeedback would not make a difference). To obviate these concerns, researchers need to employ blinded control conditions or/and transfer tasks. Gold-standard control conditions involve delivering participants feedback that should be unaffected by meditation (e.g. activations from another brain area, from another subject, or reversed activation) (Sorger et al., 2019; Thibault et al., 2016). A weaker control condition is mindfulness-as-usual, which is effective for examining general neurofeedback mechanisms, and neurofeedback benefits, but not target-specific mechanisms (Ros et al., 2020). Transfer tasks involve asking participants to meditate but removing the influence of neurofeedback- and one can examine differences in transfer tasks assessed before and after neurofeedback. Notably, the fMRI studies that employed sham controls or transfer tasks did not find significant differential evidence for target engagement (Kim et al., 2019; Kirlic et al., 2022). EEG studies did not employ transfer tasks, and only one study employed sham (Brandmeyer & Delorme 2020). Brandmeyer and Delorme (2020) found evidence of increased target engagement of frontal midline theta in mbNF, while the sham group showed no significant changes.

Comparisons of EEG-neurofeedback to mindfulness-as-usual also resulted in improved target engagement (Hinterberger & Fürnrohr, 2016; Salminen et al., 2023). In summary, there is not currently evidence from the strongest designs supporting mbNF-specific mechanisms of DMN activation control, while there are some indications of control over frontal midline theta.

### 4.4 State Mindfulness

It is critical to identify whether neural feedback can engage the proposed target mechanism, but this is insufficient if it does not yield greater mindfulness and associated mental health benefits. Ideally, target engagement also leads to increases in state mindfulness, or deeper mindfulness during practice. There is only limited evidence in our included studies for increased state mindfulness (Hinterberger & Fürnrohr, 2016; Kim et al., 2019; Kirlic et al., 2022; Zhang et al., 2023 c.f., Kim et al., 2019, Prestel et al., 2019). One concern is that monitoring of the feedback may cause distraction during meditation. For this reason, some studies provided feedback intermittently after blocks of meditation (Pamplona et al., 2020, 2023), or allowed practitioners to close their eyes during meditation (van Lutterveld et al, 2017). The studies mostly used visual feedback, which may be distracting. Future research could examine the impact of design choices on state mindfulness during mbNF, including visual/auditory modality, continuous vs intermittent feedback, etc. It may also be useful to collect data throughout the course of mbNF to assess inattention.

### 4.5 Behavioral Outcomes

Of note, mbNF has often been proposed to enhance mindfulness acquisition (Brandmeyer & Delorme, 2013). A prime motivation for many of the studies reviewed was the possibility of beneficial outcomes in cognition and affect. An fMRI study observed some improvements in reaction times on a cognitive task beyond a control condition (Pamplona et al., 2020), but it wasn’t maintained at follow-up. An EEG study identified memory improvements but not RT improvements (Brandmeyer & Delorme, 2020). Another fMRI study tested perceived stress and negative affect, and didn’t observe any improvements (Kirlic et al., 2022), whereas an EEG study identified improvements beyond a waitlist control (Dunham et al., 2018). Two fMRI studies observed decreases in auditory hallucinations (Bauer et al., 2020; Okano et al., 2020), a striking finding with implications for deleterious psychosis symptoms, however the findings were extracted from two protocols on the same sample. Of course, studies may have measured cognitive and affective outcomes but not reported them (many EEG studies did not report behavioral outcomes). Preregistration of measures and analyses was scarce. Overall, there is limited existing evidence for mindfulness-based neurofeedback benefits in terms of behavioral or clinical outcomes.

### 4.6 Clinical Relevance

Clinical populations may benefit from adaptations of mindfulness instruction. Individuals with histories of trauma may experience traumatic re-experiencing and distress due to meditation (Treleaven, 2018; Zhu et al., 2019). Ruminative individuals with a tendency to engage in repetitive negative thoughts may particularly have trouble learning meditation (Alleva et al., 2014; Crane & Williams, 2010; Hilton et al., 2017). It may be especially helpful for these clinical populations to have scaffolds while they meditate. Mindfulness-based NF may be such a scaffold, providing an engaging external locus of attention plus the same essential components of mindfulness - redirection of attention and non-judgement. Studies on mbNF included here involved clinical participants (Bauer et al., 2020, Okano et al., 2020, Zhang et al., 2023) but none involved healthy control groups. Future studies should assess directly whether the benefits of mbNF are more pronounced in clinical groups. Of course, mbNF should not be considered a replacement for more traditional mindfulness training (e.g.,with in-person teaching). There is a rich psychotherapeutic literature on developing mindfulness adaptations for clinical groups (e.g. mindfulness-based cognitive therapy; Segal et al., 2004) and acceptance and commitment therapy; (Hayes et al., 1999), and mbNF requires more validation before joining these frontline treatments.

### 4.7 Summary

This systematic review of mindfulness meditation concurrent with EEG or fMRI neurofeedback suggests that participants can learn to downregulate the DMN and increase power in the theta band. However, the lack of adequate control conditions limits mechanistic assertions. In addition, the downstream benefits of mindfulness-based neurofeedback require systematic examination. There is evidence for the feasibility of neurofeedback with clinical populations, and future work should directly compare the effects of mbNF between clinical and non-clinical populations.

### 4.7 Limitations

Our conclusions should be tempered in light of the heterogeneity of the studies. Targets, outcomes, and sample characteristics varied widely across the studies. These differences are well-known to affect neural outcomes (e.g. neuromaturation in adolescents, Fan et al., 2021; Norbom et al., 2021). Mindfulness training may be more effective for reducing psychological distress than for improving cognitive function ((Gill et al., 2020; Whitfield et al., 2022), and it is unclear whether this applies to mbNF. Reporting was also variable, which we assessed using the CRED-NF checklist (Ros et al., 2020). The vast majority of studies lacked blinded control conditions, reported brain target engagement as a single outcome instead of comprehensively, and did not engage in open science practices. In the future, full reporting of targets and outcomes could help identify why some studies may see effects and others do not, and it could lead to possible quantitative synthesis of effects.

Another limitation is the scope of the review. We chose not to review all meditation based neurofeedback, restricting our selection to studies that employed mindfulness practices. There are multiple families of meditations, including attentional, constructive and deconstructive practices (Dahl et al., 2015). The studies included here involved attentional practices. Future work should examine the effects of neurofeedback on other practices, perhaps targeting different brain processes.

### 4.8 Future Directions

To study the mechanisms of mbNF and associated effects, the field would benefit from adopting best practices. Chief among these may be a confirmatory-exploratory distinction. Exploratory studies may examine multiple targets, multiple modalities of feedback, qualitative as well as quantitative feedback – all with the aim of establishing preliminary hypotheses about neural targets. These studies are necessary and important given the nascency of the field. Three studies reviewed provide a sound roadmap for this type of work (Garrison et al., 2013a b, Van Lutterveld et al., 2017). One innovation in particular is working with experienced meditators, who have detailed awareness of mental phenomena during meditation. Another innovation is developing individualized targets - one method could be monitoring neural data during meditation for a given participant, with self-report probes (experience samples), and then in a subsequent task delivering feedback that was trained on that initial period. This personalization may be more effective than using ‘one-size-fits-all’ brain signals (Brandmeyer & Reggente, 2023).

It is critically important, however, to build on this work using RCTs with carefully designed sham control conditions. Such confirmatory work, through tests of clear and a priori hypotheses, can help the field evaluate whether participants learn to modulate a neural signal, and whether it leads to higher state mindfulness and positive mental health or cognitive outcomes. Sham or alternative ROI controls are preferred, given their ability to control for effects of placebo as well as feedback monitoring, but mindfulness-as-usual controls are useful and easier to implement. Researchers may even choose to examine different dosages of mbNF (Bloom et al., 2023). Clinical trial registration and/or preregistration is useful, and when deviations emerge as they always do during empirical research, they should be reported. As mentioned previously, the CRED-NF checklist should be used for standardized reporting.

A final aim is real-world translation. In contrast to fMRI, which is costly and largely only accessible via academic medical centers, there is burgeoning interest in consumer-grade EEG tools like MUSE (Hashemi et al., 2016; Sawangjai et al., 2019), which are relatively cheap (∼$250) and easy to use. We believe that this interest should be tempered given the limited knowledge base in lab settings. EEG tools like MUSE may rely on the potent influence of *neurosuggestion* (Schönenberg et al., 2017), which is a cultural emphasis and trust in Western society for neuroscientific technology. Speculatively, *neurosuggestion* effects may not be sustainable in supporting a habit of meditation, and may obscure the self-insight that comes with meditation (Vago & David, 2012).

## Data and Code Availability

N/A

## CRediT Statement

INT contributed conceptualization, methodology, investigation, writing-original draft, writing- review & editing, visualization and project administration. KDG contributed investigation, writing-original draft, writing- review & editing, and visualization. EW & ZB contributed investigation, methodology and writing- review & editing. PAB, NK, DP, JZ, CCB contributed review & editing. SWG contributed supervision, funding acquisition. RPA contributed supervision, review & editing, funding acquisition.

## Ethics Statement

This article does not contain original research.

