## Supplement for "Mindfulness-based Neurofeedback: A Systematic Review of EEG and fMRI studies"

| **Section and Topic** | **Item #** | **Checklist item** | **Location where item is reported** |
| --- | --- | --- | --- |
| **TITLE** | | |  |
| Title | 1 | Identify the report as a literature review. | Pg. 1 |
| **ABSTRACT** | | |  |
| Abstract | 2 | Provide a structured summary including, as applicable: background; objectives; data sources; study eligibility criteria, participants, and interventions; study appraisal and synthesis methods; results; limitations; conclusions and implications of key findings.  See the [PRISMA 2020 for Abstracts checklist](http://www.prisma-statement.org/Extensions/Abstracts.aspx) for the complete list. | Pg. 2 |
| **INTRODUCTION** | | |  |
| Rationale | 3 | Describe the rationale for the review in the context of existing knowledge, i.e., what is already known about your topic. | Pg. 3-7 |
| Objectives | 4 | Provide an explicit statement of the objective(s) or question(s) the review addresses with reference to participants, interventions, comparisons, outcomes, and study design (PICOS). | Pg. 6, Par. 2 |
| **METHODS** | | |  |
| Eligibility criteria | 5 | Specify the inclusion and exclusion criteria for the review and how studies were grouped for the syntheses with study characteristics (e.g., PICOS, length of follow-up) and report characteristics (e.g., years considered, language, publication status) used as criteria for eligibility, giving rationale. | Pg. 7, Par. 2 |
| Information sources | 6 | Specify all databases, registers, websites, organisations, reference lists and other sources searched or consulted to identify studies. Specify the date when each source was last searched or consulted. | Pg. 8, Par. 1 |
| Search strategy | 7 | Present the full search strategies for all databases, registers and websites, including any filters and limits used. | Pg. 8, Par. 1 |
| Selection process | 8 | State the process for selecting studies (i.e., screening, eligibility).  Specify the methods used to decide whether a study met the inclusion criteria of the review, including how many reviewers screened each record and each report retrieved, whether they worked independently, and if applicable, details of automation tools used in the process. | Pg. 8, Par. 2 |
| Study risk of bias assessment | 11 | Specify the methods used to assess risk of bias in the included studies, including details of the tool(s) used, how many reviewers assessed each study and whether they worked independently, and if applicable, details of automation tools used in the process. | Pg 8. , par. 4 |
| **RESULTS** | | |  |
| Study selection | 16a | Describe the results of the search and selection process, from the number of records identified in the search to the number of studies included in the review, ideally using a flow diagram. | Pg. 9 Par. 2 |
|  | 16b | Cite studies that might appear to meet the inclusion criteria, but which were excluded, and explain why they were excluded. | NA |
| Study characteristics | 17 | Cite each included study and present its characteristics (e.g., study size, PICOS, follow-up period). | Tables 1 and 2 |
| Risk of bias in studies | 18 | Present assessments of risk of bias for each included study. | Pg. 18, par.3, pg. 27 |
| Results of individual studies | 19 | For all outcomes, present, for each study: (a) summary statistics for each group (where appropriate) and (b) an effect estimate and its precision (e.g. confidence/credible interval), ideally using structured tables or plots. | NA |
| **DISCUSSION** | | |  |
| Discussion | 23a | Provide a general interpretation of the results in the context of other evidence. | Pg. 28-33 |
|  | 23b | Discuss any limitations of the evidence included in the review. | Pg. 33, par. 2 |
|  | 23c | Discuss any limitations of the review processes used. | Pg. 33 par. 2 |
|  | 23d | Discuss implications of the results for practice, policy, and future research. | Pg. 33-34 |
| **OTHER INFORMATION** | | |  |
| Registration and protocol | 24a | Provide registration information for the review, including register name and registration number, or state that the review was not registered. | NA |
|  | 24b | Indicate where the review protocol can be accessed, or state that a protocol was not prepared. | NA |
|  | 24c | Describe and explain any amendments to information provided at registration or in the protocol. | NA |
| Support | 25 | Describe sources of financial or non-financial support for the review, and the role of the funders or sponsors in the review. | Pg. 34-35 |
| Competing interests | 26 | Declare any competing interests of review authors. | Pg. 34-35 |
| Availability of data, code, and other materials | 27 | Report which of the following are publicly available and where they can be found: template data collection forms; data extracted from included studies; data used for all analyses; analytic code; any other materials used in the review. | NA |
