## Supplement 2 for "Mindfulness-based Neurofeedback: A Systematic Review of EEG and fMRI studies"

**Table S1: CRED-NF items, full**

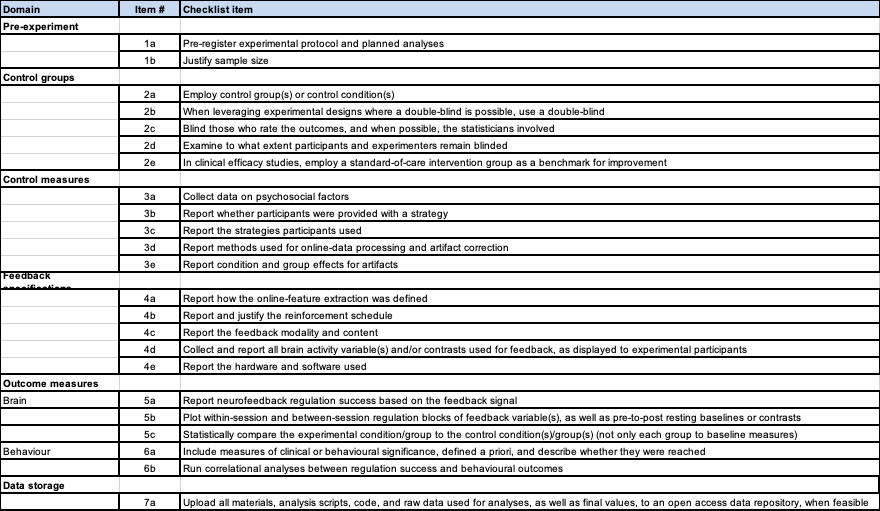

**Table S2: fMRI CRED-NF coding**

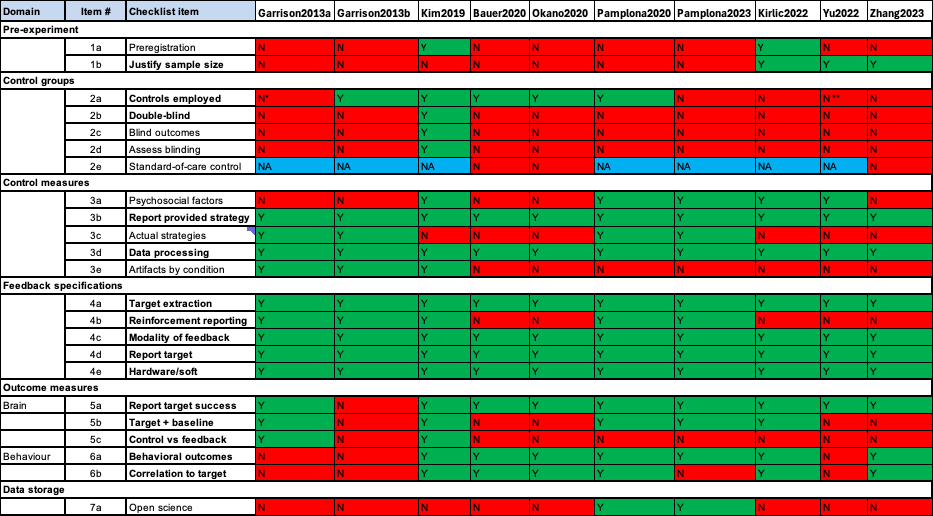

Bold items reflect essential checklist items. Please note that Garrison2013a and Garrison 2013b used same protocol, Bauer2020 and Okano2020 used same sample, Pamplona2020 and Pamplona2023 used same sample, and Kirlic2022 and Yu2022 used same sample. This means we coded ‘Y’ for those studies if either manuscript reported a methodological detail (but not for outcomes). NA: not applicable, in the case of non-clinical samples, there is no ‘standard-of-care’. Note that Kirlic2022 and Yu2022 filled out checklists, which we re-coded where necessary for this review. * Garrison2013a reported non-meditators as a control but they also received mbNF. ** Yu2022 reported controls in their checklist, but their controls are not between-subject or within-subject, instead they are just baseline resting-state.

**Table S3: EEG CRED-NF coding**

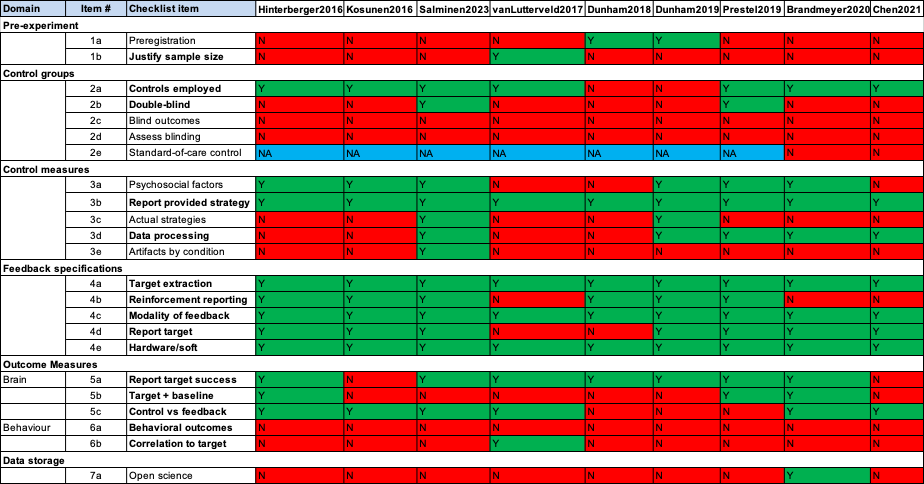

Bold items reflect essential checklist items. Hinterberger2016: Hinterberger & Funrohr 2016, Kosunen2016: Kosunen et al., 2016, Salminen2023: Salminen et al., 2023, vanLutterveld2017: van Lutterveld et al., 2017, Dunham2018: Dunham et al., 2018, Dunham2019: Dunham et al., 2019, Prestel2019: Prestel et al., 2019, Brandmeyer2020: Brandmeyer & Delorme, 2020, Chen: Chen et al., 2021 etc.

**Table S3: fMRI Qualitative Ratings**

| Garrison, Scheinost, et al., 2013 | Exp 1: After each run, participants rated how well the graph corresponded with their moment-to-moment subjective experience, briefly described this rating, and rated how well they were able to follow instructions.  Exp 2: After each run, participants described their experience during focused attention meditation, rated how well the graph corresponded with experiencing during meditation (for both offline and real-time feedback), and reported what strategy they used to decrease the feedback graph. |
| --- | --- |
| Garrison, Santoyo, et al., 2013 | Same as Exp 2 in Garrison, Scheinost, et al., 2013 |
| Kim et al., 2019 | Not reported |
| Bauer et al., 2020 | Not reported |
| Okano et al., 2020 | STG feedback: Participants rated how well they were able to attend to the voices after each listen block, and how well they were able to ignore all sounds after each ignored block.  SMC feedback: After each block, participants rated how vigorously they moved their fingers. |
| Pamplona et al., 2020 | After each NF run, participants rated their control over the thermometer, how difficult it was to control the thermometer, and their concentration level. |
| Pamplona et al., 2023 | After each NF run, participants rated what strategy they used and their concentration level. |
| Kirlic et al., 2022 | After each run, participants reported how well they were able to follow instructions on the screen, how easy they found it to focus on breath, how much their mind wandered, how easy it was to mentally decide whether or not words described them, how easy it was to clear their mind while resting, and how they felt.  After each NF run, participants reported how well the blue bar corresponded with their experience of breath focus, and how well the red bar corresponded with their experience of mind wandering. |
| Yu et al., 2022 | Same as Kirlic et al., 2022 |
| Zhang et al., 2023 | After each run, participants confirmed that they had used mental noting during NF |

**Table S4: EEG Qualitative Ratings**

| Hinterberger & Fürnrohr, 2016 | Participants completed a feedback questionnaire after each interventional condition, rating their physical sensation, emotional condition, mental state, experience, motivation, duration, and relevance. Participants were also asked, “Could you notice a connection between yourself and the audiovisual perceptions?” |
| --- | --- |
| Kosunen et al., 2016 | After each experiment condition, participants completed the ITC-Sense of Presence Inventory and a meditation depth questionnaire measuring five factors: Hindrance, Relaxation, Personal Self, Transpersonal Qualities, and Transpersonal Self. |
| Salminen et al., 2023 | After each experiment condition, participants completed the ITC-Sense of Presence Inventory and the 6-item factor ‘Hindrances’ from a meditation depth questionnaire. |
| van Lutterveld et al., 2017 | Participants were asked which direction of the graph they associated with their subjective experience of effortless awareness, and rated their confidence in their response. |
| Dunham et al., 2018 | Not reported |
| Dunham et al., 2019 | After session 2 of each learning day, participants rated their perceptions that the following cognitive states were associated with reductions in the BIS values: widening the visual field, decreasing effort, attention to space, and relaxed alertness. |
| Prestel et al., 2019 | Participants completed a semi-structured interview immediately after each NF session, reporting on their perceived success in changing the signal, which strategies they applied during each block, difficulties or distractions, and an overall evaluation of the session. |
| Brandmeyer & Delorme, 2020 | Participants reported on whether they were able to successfully implement one of the learned strategies after each day of NF. |
| Chen et al., 2021 | Not reported |

**Figure S1: Sample sizes for studies**

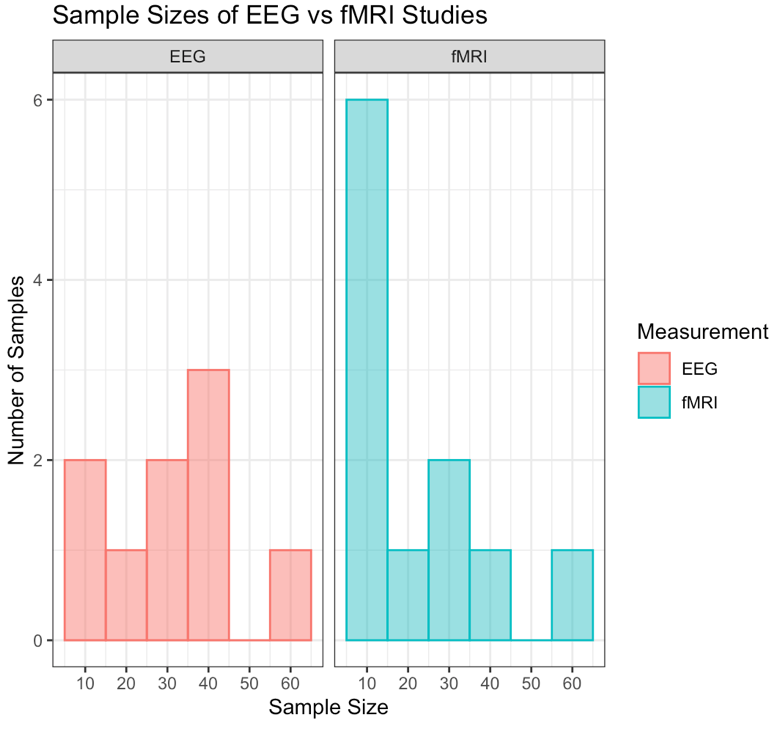

**Figure S1.** Side by side histograms comparing the sample sizes of EEG studies and fMRI studies. Duplicate samples are included.

**Figure S2: Sample sizes for unique samples**

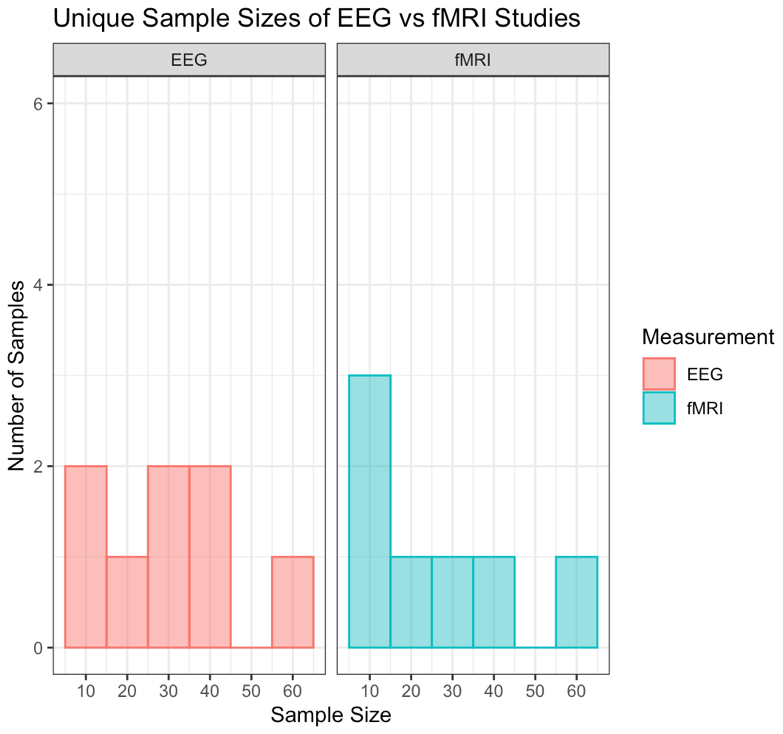

**Figure S2.** Side by side histograms comparing the unique sample sizes of EEG studies and fMRI studies. Duplicate samples are not included. In cases where duplicate samples have different sample sizes, the higher sample size was included.

**Figure S3: Control conditions by study**

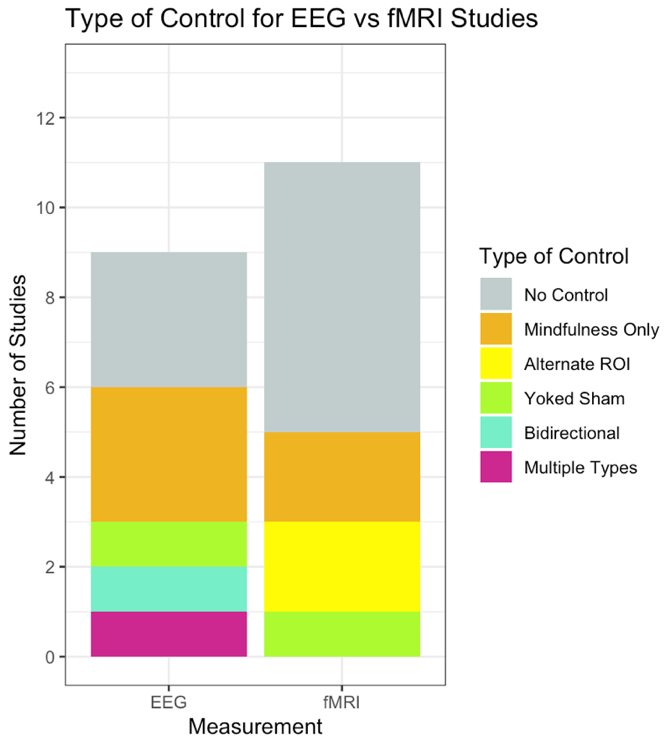

**Figure S3.** Stacked bar chart showing the types of control conditions in EEG and fMRI studies. Studies that included more than one control condition are counted as “multiple types.” There are more controlled EEG studies, largely driven by ‘mindfulness only’ controls. On the other hand, there are alternate ROI controls in the fMRI studies which are not present in EEG.

**Figure S4: Control conditions overall**

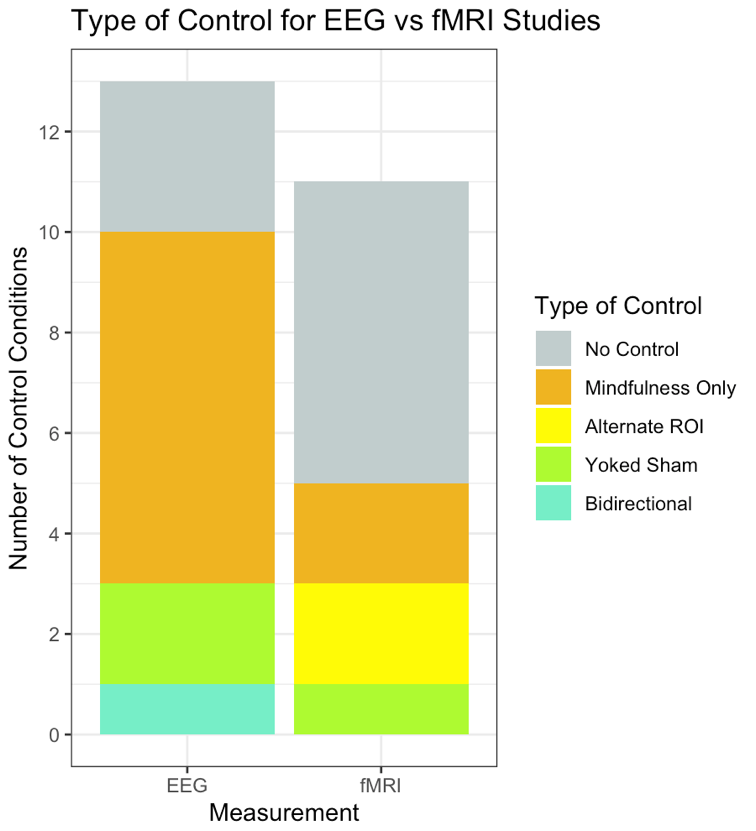

**Figure S4.** Stacked bar chart showing the types of control conditions in EEG and fMRI studies. Studies that included more than one control condition count towards each type of control included.

**Figure S5: Types of control conditions**

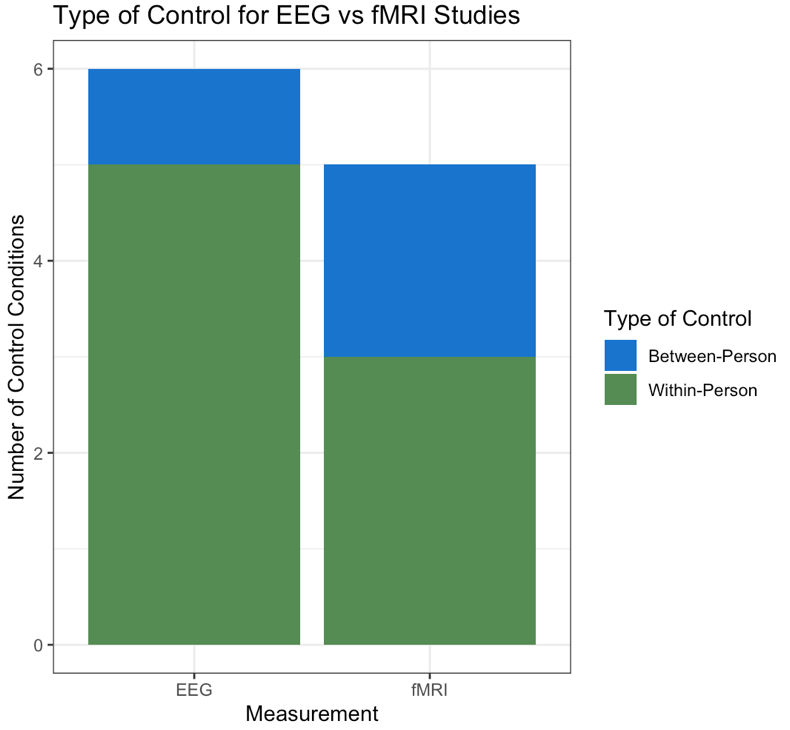

**Figure S5.** For studies with control conditions, what are the quantities of between-person and within-person controls for EEG and fMRI studies. EEG studies are predominantly within-person, whereas fMRI involved two between-person controls.
